# Aggregation Dynamics of a 150 kDa Aβ42 Oligomer: Insights from Cryo Electron Microscopy and Multimodal Analysis

**DOI:** 10.1101/2024.07.30.605873

**Authors:** S. Shirin Kamalaldinezabadi, Jens O. Watzlawik, Terrone L. Rosenberry, Anant K. Paravastu, Scott M. Stagg

## Abstract

Protein misfolding is a widespread phenomenon that can result in the formation of protein aggregates, which are markers of various disease states, including Alzheimer’s disease (AD). In AD, amyloid beta (Aβ) peptides, particularly Aβ40 and Aβ42, are key players in the disease’s progression, as they aggregate to form amyloid plaques and contribute to neuronal toxicity. Recent research has shifted attention from solely Aβ fibrils to also include Aβ protofibrils and oligomers as potentially critical pathogenic agents. Particularly, oligomers demonstrate greater toxicity compared to other Aβ specie. Hence, there is an increased interest in studying the correlation between toxicity and their structure and aggregation pathway. The present study investigates the aggregation of a 150 kDa Aβ42 oligomer that does not lead to fibril formation over time. Using negative stain transmission electron microscopy (TEM), size exclusion chromatography (SEC), dynamic light scattering (DLS), and cryo-electron microscopy (cryo-EM), we demonstrate that 150 kDa Aβ42 oligomers form higher-order string-like assemblies over time. The strings are unique from the classical Aβ fibril structures. The significance of our work lies in elucidating molecular behavior of a novel non-fibrillar form of Aβ42 aggregate.

## II. Introduction

Protein misfolding is a common phenomenon in nature that may result in the formation of protein aggregates known to be the hallmark of various diseases, including Alzheimer’s disease (AD) (1–3). In AD, amyloid beta (Aβ) peptides, particularly Aβ40 and Aβ42, play central roles, aggregating, forming amyloid plaques, and contributing to neuronal toxicity (4–8). Recent discoveries shifted the focus from Aβ fibrils as the sole pathogenic agents to Aβ protofibrils and oligomers. In this work, we studied the aggregation behavior of 150 kDa (32-mer) oligomers of the Alzheimer’s amyloid-β42 (Aβ42) peptide. Oligomers represent an important aspect of the complex protein aggregation phenomena that underly Alzheimer’s disease (9–14). We do not know why oligomers are limited in size to ∼100’s of molecules while thousands of Aβ molecules can assemble to form amyloid fibrils. This observation is mysterious when we consider that oligomers and fibrils share the same β-sheet structural motif (15–23); the distinct behavior of oligomers and fibrils is likely related to their differing β-sheet architectures. Aβ fibrils are composed of β-strands arranged perpendicular to the fibril axis (“cross-β” motif) with consistent inter-strand alignments (usually in-register parallel) (24). Generally, a planar β-sheet, irrespective of size, would be expected to recruit additional β-strand molecules into itself due to dangling hydrogen bond donors and acceptors, thereby growing in length. In contrast, our recent work using solid-state NMR measurements on 150 kDa Aβ42 oligomers have revelated a more complex arrangement of β-strands, which include the coexistence of both parallel and antiparallel β-sheets (15). Here, we show that 150 kDa Aβ42 oligomers do exhibit a degree of further aggregation, but this behavior is distinct from fibril formation.

We are interested in oligomeric Aβ aggregation pathways because oligomers are believed to play a special role in pathology (10–14). Aβ can form a diverse array of non-fibrillar structures, broadly classified as oligomers and protofibrils, and the relationships between structure, aggregation mechanisms, and pathology remain poorly understood (25). Although the amyloid cascade hypothesis proposes that the aggregation of Aβ peptides in fibrillar form contributes to neuronal toxicity and cognitive decline in AD (10–14,26), recent research has indicated that the degree of dementia demonstrates a stronger correlation with the concentration of soluble Aβ species rather than the plaque count (27). In support of this statement, inhibition of Aβ fibril formation does not always reduce Aβ toxicity in cultured neurons, while oligomers and protofibrils can alter neuronal function and cause cell death (28). Oligomers could potentially be highly toxic due to their hydrophobic interactions with lipid membranes (29) (30), and their high diffusion rates, allowing abnormal interactions with various cellular components like phospholipid bilayers, receptors, RNAs, proteins, and metabolites (24). Aβ oligomers disrupt phospholipid bilayers, causing membrane curvature and discontinuity (30)(31)(11), and can form pore-like structures contributing to membrane disruption (28). There appears to be a consensus among several studies that oligomers/protofibrils and not the monomers or fibrils are the most toxic Aβ species (30,32–35). Additionally, Several different studies demonstrated that Aβ oligomers added to cell culture media are heterogeneous and are likely to change their assembly state during an experiment (36–40). Despite their potential importance, the distinct aggregation mechanisms of oligomers remain uncertain.

Although studies of oligomers can be very challenging, a 150 kDa Aβ42 oligomer has exhibited sufficient structural and homogeneity for characterization. In 2007, Rangachari *et al*. reported that 150 kDa Aβ42 oligomers are formed when monomeric Aβ42 undergoes conversion into predominantly β-structured conformations in 2 mM sodium dodecyl sulphate (SDS), which do not proceed to form fibrils (41). In 2009, Moore *et al*. generated soluble Aβ42 oligomers through incubation of synthetic peptides in dilute SDS followed by dialysis to remove SDS, resulting in an oligomer with a mass of 150 kDa rich in β-sheet (42). Tay *et al*. (2013) used NMR to show that the intermolecular distances for the β-sheets in 150 kDa Aβ42 oligomers were inconsistent with the in-register parallel β-sheet structure observed in fibrils. The 150 kDa Aβ42 oligomers also do not exhibit thioflavin T fluorescence, likely due to their distinct intermolecular β-strands (43). In 2015, Huang *et al*. performed solid-state NMR studies on the 150 kDa Aβ42 oligomer, which suggested that the C-terminal β-strand was arranged as an antiparallel β-sheet (44). In contrast, most Aβ42 fibril structures are known to have parallel β-sheets. In 2020, Gao *et al*. conducted similar NMR studies and discovered that residues 11-24 of Aβ42 oligomers form an out-of-register parallel β-sheet in the same structure (15). Most fibrillar structures are reported to have in-register parallel β-sheets. As such, the coexistence of parallel and antiparallel β-sheets may distinguish non-fibrillar forms from fibrillar structures (15). Altogether, these data indicate that the 150 kDa Aβ42 oligomers are off pathway to fibril formation due to their distinct β-sheets arrangement.

Since the toxicity of Aβ aggregates is structure-dependent (15), a deeper and more detailed understanding of Aβ oligomer assembly is crucial for comprehending AD pathology at a molecular level. Our purpose is to characterize an aggregation pathway that is formed by Aβ42 oligomers without further assembly into fibrils. The results of our study using negative stain TEM, size exclusion chromatography, and dynamic light scattering suggest that the 150 kDa Aβ42 oligomers form higher-order string-like assemblies, and the “strings” remain stable for weeks. We study how the addition of NaCl or SDS can affect the formed strings and demonstrate monomer addition can disrupt the strings formed by these oligomers. We further demonstrate that the oligomeric “strings” are off-pathway to fibrillar formation. Specifically, we observed that these strings are distinctly different from classic Aβ fibrils due to differences in their assembly; strings are assembled from 150 kDa Aβ42 oligomers, while fibrils are assembled from Aβ42 monomers. We also used cryo-EM and 2D classification to elucidate the assembly of oligomers into strings. This type of self-association is not highly ordered like what is observed in fibrils, which can explain why the strings are limited in length. Altogether, our data demonstrate that the 150 kDa Aβ42 oligomers organize into novel higher-order string-like structures over time that are off the pathway to fibril formation.

## III. Materials and Methods

### Aβ42 peptide synthesis, monomer isolation and oligomer preparation

Aβ42 peptides (sequence DAEFR HDSGY EVHHQ KLVFF AEDVG SNKGA IIGLM VGGVV IA) were purchased from the peptide synthesis core facility at the Mayo Clinic (Rochester, MN), and the purity was determined by MALDI-mass spectrometry to be > 90%. The monomer sample was isolated in 20 mM sodium phosphate pH 7.5 (43,44) and flash-frozen in liquid nitrogen. 150 kDa Aβ42 oligomer preparation was started by thawing 1.5 mL aliquots of monomeric Aβ42 solutions at room temperature (25 °C). To initiate formation of 2-4mer oligomers, SDS detergent was added to a final concentration of 4 mM (Aβ42 monomer concentration ∼100 µM). Samples were then incubated at room temperature for 48 hours without agitation followed by removal of the detergent to induce conversion of 2-4mer oligomers to 150 kDa (∼32mer) oligomers. Detergent was removed either by dialysis alone, or with dialysis and BioBeads depending on the experiment described in the results. For dialysis alone, each sample was transferred to dialysis tubing (Thermo Scientific SnakeSkin Dialysis Tubing, 7000 MWCO, cat# 68700) and dialyzed against 500-700 mL of 10 mM sodium phosphate pH 7.5 (dialysis buffer) at room temperature on a stir plate. For the experiments where ionic strength was varied, the dialysis buffer was supplemented with sodium phosphate and sodium chloride as described in the Results. The dialysis buffer was exchanged at least 5 times in 48 hours. For dialysis with BioBeads, the sample was dialyzed for only 2 hours against one liter of dialysis buffer followed by addition of 200 mg Bio-Beads SM2 (cat# 1523920) per mL of Aβ42 sample. The sample mixed with BioBeads was rotated for two hours at room temperature in a 15 mL Corning tube using a VWR Multimix Tube Rotator Mixer (cat# 13916-822). After 2 hours of contact with BioBeads, the tube was spun at 3200 × g several times to precipitate the beads. Following SDS removal samples were injected onto a Superose 6-increase 10/300 GL column (Cytiva) pre-equilibrated with dialysis buffer. The flow rate was kept at 0.2 mL/min during the run and 0.5 mL fractions were collected. The fractions associated with 150 kDa oligomer were retained for further analysis.

### Negative-stain TEM visualization of Aβ42 samples

Carbon film 400 mesh, Cu grids (electron microscopy sciences CF400-CU) were plasma cleaned with a Solarus 950, Gatan Advanced Plasma System for 20 seconds, within an hour before applying the sample. 4 µL of Aβ42 oligomer sample was applied at the top of each grid, and after 2 minutes, the non-adhered sample was blotted off using a Whatman filter paper (cat# 1001090). The grid was then washed twice with 4 µL of filtered autoclaved nano-pure water Thereafter, 4 µL of 2 % uranyl acetate solution was added to the surface. After two minutes contact with stain, the excess uranyl acetate was blotted off. The grids were visualized using a Hitachi 7800 Transmission Electron Microscope at FSU biological imaging resource (BSIR) facility.

### Sample preparation for performing size exclusion chromatography at different timepoints after SDS removal

Aβ42 oligomer sample was prepared using dialysis and BioBeads as described above. For each SEC run 500 μL of sample was injected onto a Superose 6 increase 10/300 GL column pre-equilibrated with dialysis buffer. The first SEC run was performed immediately after SDS removal. The remaining of the sample was kept at 4 degrees and the second SEC run was performed 48 hours later.

### Dynamic light scattering

Aβ42 oligomer sample was prepared by dialysis and BioBeads, but no SEC was performed on the oligomer sample prep after removal of SDS. After preparation, the sample was kept at 4 °C for storage. The DLS measurements were performed using a DynaPro NanoStar instrument (WYATT Technology) using a 1 µL black-walled quartz micro-cuvette. The temperature was set at 25 °C. For measurements, the DLS acquisition time was 2 seconds, set number of DLS acquisitions was 10, and for each sample the same loop was repeated 3 times. 5 µL of sample was added to the cuvette each time for these measurements and the measurement was repeated at least 2 times by cleaning the cuvette and adding another 5 µL of sample.

### Sample preparation for studying the effect of addition of salt, SDS and monomer to Aβ42 oligomer sample

Aβ42 oligomer sample was prepared by SDS removal with dialysis alone for studying the effect of salt and SDS on sample. High concentration stocks of these solutions were made and filtered. Thereafter, for 6 mM salt: 4 µL of Aβ42 oligomer sample was mixed with 1µL of 30 mM NaCl and for 6 mM SDS: 4 µL of Aβ42 oligomer sample was mixed with 1µL of 30 mM SDS. After 1-2 minutes, 4 µL of each tube was taken and used to prepare negatively stained grids.

For studying the effect of monomer addition to Aβ42 oligomer, oligomer sample was prepared using dialysis and BioBeads. After SEC, the oligomer peak fraction was kept at 4 °C. After 6 days, the sample was used to take negative stain images using TEM. Thereafter, a 1:1 v/v ratio of oligomer sample to 178 µM monomer was prepared. After a 1 min incubation at room temperature, this mix was used to prepare negatively stained samples of monomer plus oligomer. This procedure was repeated on the original oligomer sample 2 days later to confirm these results.

### Cryo-EM of strings

The Aβ42 oligomers were prepared using dialysis alone and concentrated to 800 μM. Thereafter, the sample was diluted 1:1 with dialysis buffer and frozen using a SPT Labtech Chameleon at New York Structural Biology Center. 6,715 micrographs were collected on this sample using a Titan Krios operated at 81,000 X magnification and 300 keV. The images were acquired on a K3 Bioquantum at a pixel size of 1.083 Å/pixel with the energy filter slit width set to 20 eV. The dose for the exposures was 60 e^-^/Å^2^. All steps of image processing were performed in CryoSPARC v4.2.1. After patch CTF estimation, the first set of particles were picked with Template Picker. The particles were inspected, and parameters were adjusted. Thereafter, several rounds of 2D classification were performed on this set of particles before getting the final 2D class averages presented here.

## IV Results

### 150 kDa Aβ42 oligomers form higher-order assemblies we call “strings”

We characterized the 150 kDa Aβ42 oligomers with negative stain TEM (**Fig. 1A**), observing that the oligomers form string-like assemblies shortly after samples preparation (**Fig. 1B**). The oligomers were prepared as previously reported: Aβ42 forms oligomers when incubated with SDS below its critical micelle concentration, and removal of SDS leads to the formation of stable 150 kDa Aβ42 oligomers (42). We hypothesized that the strings were linear assemblies of bead-shaped 150 kDa globular oligomers, so we characterized the formation of these higher-order assemblies with a series of experiments including observing their formation over time, testing the influence of ionic strength, determining if the assembly formation is reversible, and characterizing the structure with cryo-EM.

**Fig. 1.**
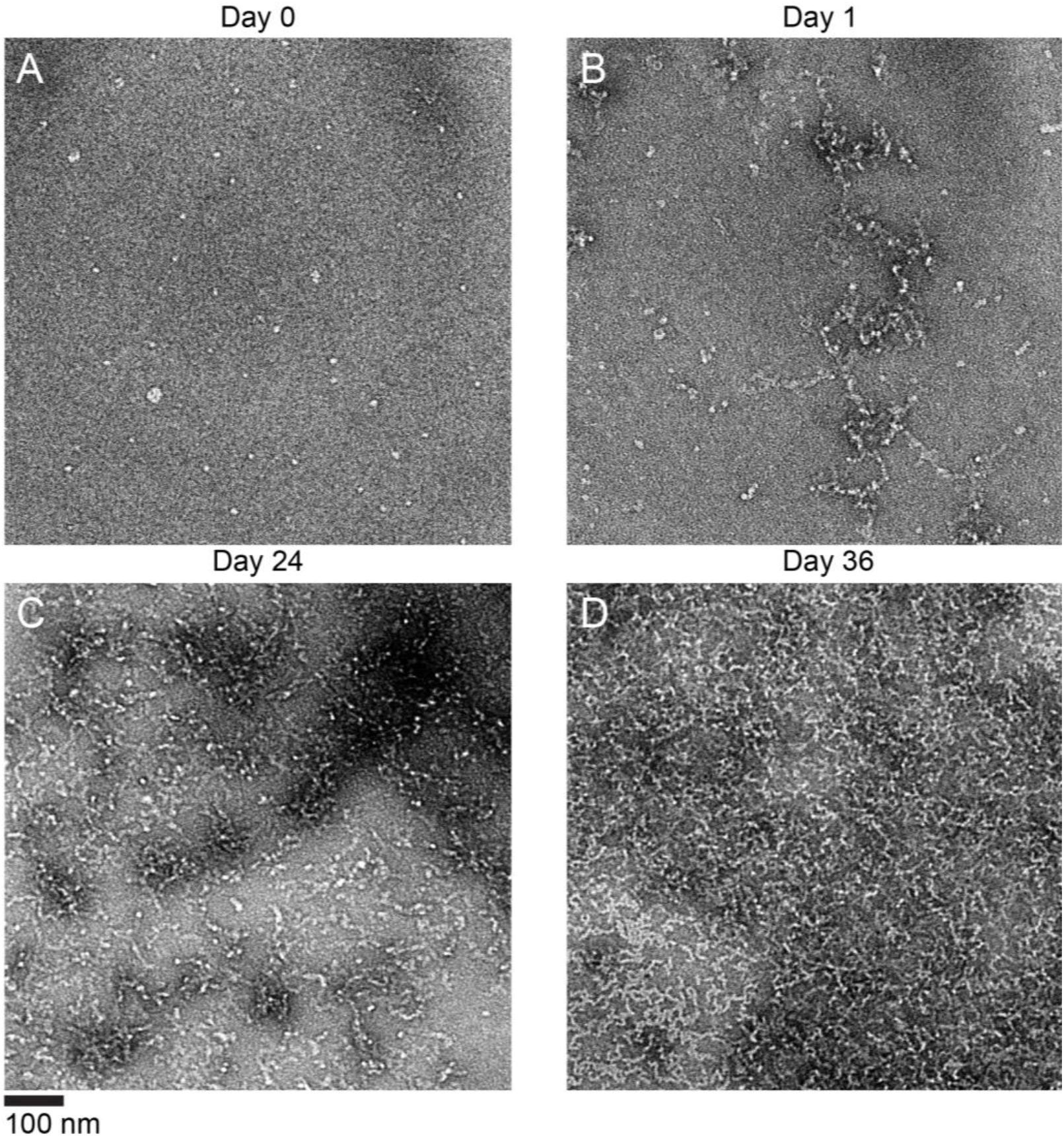
Time course of negative stain TEM images of the same Aβ42 oligomer sample prepared in low ionic strength buffer (10 mM sodium phosphate pH 7.5). Strings lengths appear to increase over time following SEC isolation of 150 kDa Aβ42 oligomers. A) Sample imaged immediately after SEC. B) Sample imaged one day after SEC. C) Sample imaged 24 days after SEC. D) Sample imaged 36 days after SEC.

First, we tested string formation in a low ionic strength buffer containing 10 mM sodium phosphate at pH 7.5. Immediately after size exclusion chromatography (SEC), the sample predominantly consisted of apparently individual globular particles that were consistent with 150 kDa Aβ42 oligomers with diameters of about 6 nm (**Fig. 1A**). When we repeated the negative stain imaging the following day, we found that the globular particles had converted to short beaded string-like structures (**Fig. 1B**). These strings varied in size from ∼12 nm to several hundred nm net-like assemblies. While the strings appeared to be linear assemblies, they were not straight. Most strings only continued a given direction ∼50 nm before changing direction with a sharp turn. Over the course of many days, the strings got longer and coalesced into patches of aggregated strings (**Fig. 1C & D**). The width of the strings remained ∼6 nm throughout the experiment.

### DLS and SEC results confirm the time-dependent formation of higher-order assemblies of Aβ42 oligomers

We hypothesized that SDS remaining in solution after dialysis might be contributing to string formation, so BioBeads were added to the sample to adsorb as much remaining SDS as possible. We then used size exclusion chromatography (SEC) and dynamic light scattering (DLS) to confirm that the higher-order assembly formation was occurring in bulk solution. The results of our experiments demonstrated that increased SDS removal was associated with increased string formation. SEC of the oligomer sample immediately after removal of SDS revealed three peaks (**Fig. 2A**). The first peak corresponds to the 150 kDa Aβ42 oligomer, and the second and third peaks correspond to partially assembled and un-assembled monomer. The SEC was repeated after 48 hours, and the results showed that the first peak shifted significantly to earlier elution volumes, indicating larger size, while the other two peaks remained at the same respective elution volumes.

**Fig. 2.**
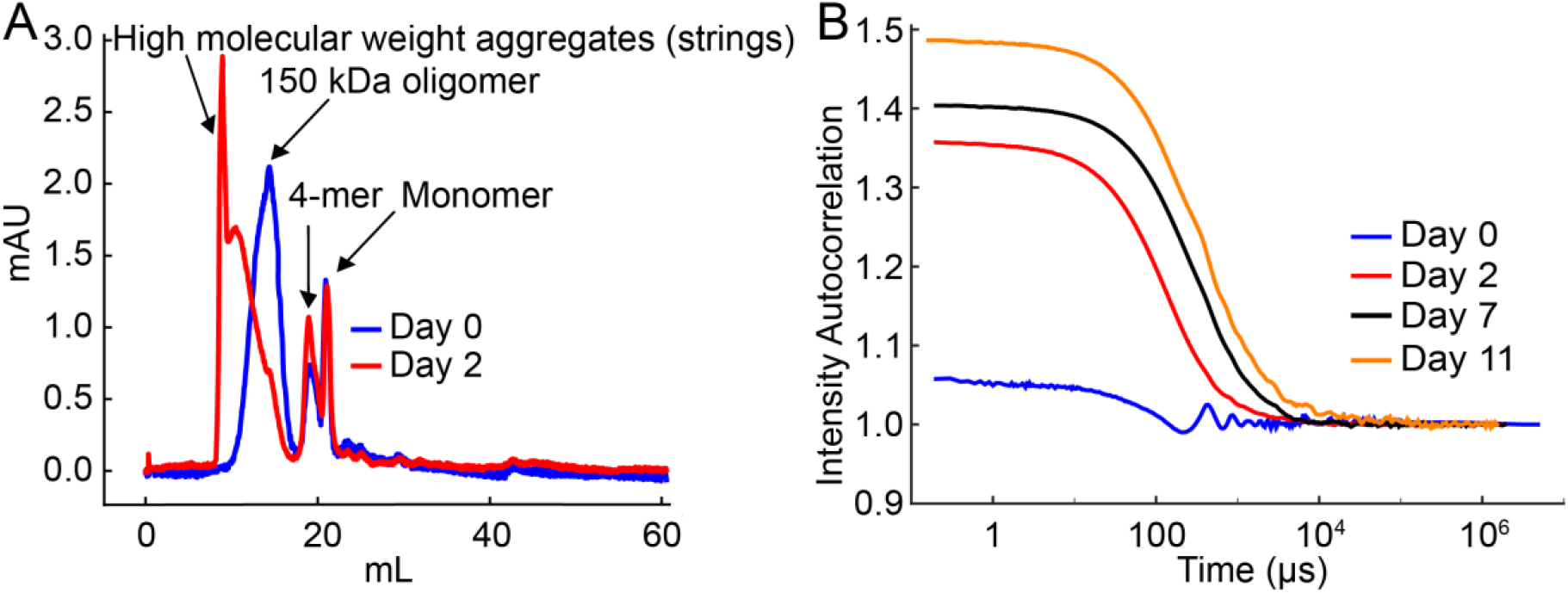
Bulk solution characterization of Aβ42 oligomers. A) SEC profile of Aβ42 sample immediately after SDS removal (blue) and two days after SDS removal (red). B) DLS autocorrelation curves at time points from 0 to 11 days.

We observed a similar result by DLS. **Fig. 2B** shows a DLS time course with autocorrelation curves collected on different days. To avoid inaccuracies of mass estimation with DLS, we only looked at the autocorrelation curves. Higher autocorrelations at earlier DLS time points are indicative of slower particle tumbling and are associated with larger particle size. The autocorrelation curves immediately after SEC were relatively flat, but the early autocorrelation increased dramatically after two days and increased steadily during the 11 days the experiment was run. Together, the negative stain imaging, SEC, and DLS experiments confirm that the individual 150 kDa Aβ42 oligomers are short-lived as individual particles, and they inevitably convert completely to strings over the course of a few days. This conversion appears to be faster when more SDS is removed.

### Doughnut shaped structures form in Aβ42 oligomer samples over time at higher ionic strength

Next, we tested whether ionic strength would influence string formation. We reasoned that if electrostatic interactions mediate string assembly, then the assembly of strings would be affected in the presence of increased salt. We repeated the time course of negative stain imaging experiments but prepared the sample in 20 mM sodium phosphate pH 7.5 supplemented with 50 mM NaCl, instead of 10 mM sodium phosphate alone. This higher ionic-strength sample exhibited a similar pattern of string formation in which the length and quantity of strings increased over time (**Fig. S1)**. However, there were a few noticeable differences. In the higher ionic-strength sample, strings were observed immediately after SEC (**Fig. S1-A**). The strings appeared to grow at a slower rate than the low ionic-strength condition, however, this is likely explained by the appearance of new doughnut-shaped assemblies (12-15 nm in diameter) that appeared after several days (**Fig. 3A**). These assemblies appear to be short strings that double back on themselves to form a ring. This interpretation is based on the observation that the doughnut wall thickness seems to be similar to string thickness (∼6 nm). The doughnuts increased in number over time (**Fig. 3B-C-D**) and were irregularly distributed on the grid, forming patches of aggregates alongside growing strings (**Fig. 3D**). To show that the doughnut formation was reproducible, we repeated the experiment three times with high salt buffer and in each case doughnuts were observed after a few days. The doughnuts were occasionally observed in samples prepared in lower ionic strength but to a lesser extent.

**Fig. 3.**
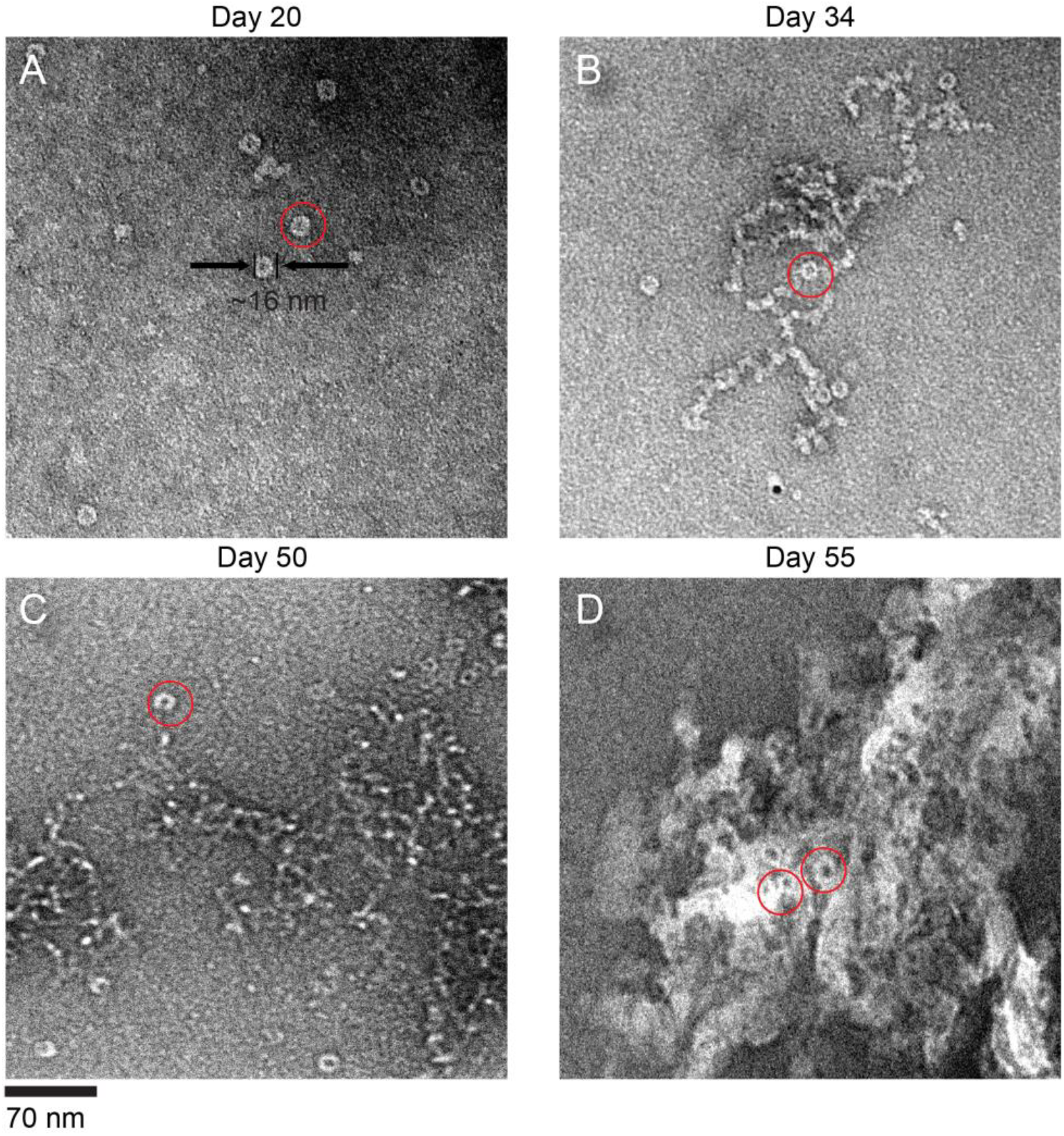
Appearance of doughnut shape structures in the Aβ42 oligomer sample prepared in buffer with higher ionic strength (20 mM sodium phosphate pH 7.5 supplemented with 50 mM NaCl). The TEM images of negatively stained sample are selected manually to exhibit the appearance of doughnuts in samples at different stages after SEC. A) Sample imaged 20 days after SEC. B) Sample imaged 34 days after SEC. C) Sample imaged 50 days after SEC. D) Sample 55 days after SEC.

### Aβ42 string formation appears to be reversible upon the addition of salt, SDS, or Aβ42 monomer

We investigated whether the string formation in Aβ42 oligomer samples was reversible. The addition of small amounts of salt (6 mM NaCl) to aged samples (24 days past SEC) composed mainly of strings (**Fig. 4A**) induced partial dissociation, leading to the formation of shorter strings and/or individual 150 kDa globular particles (**Fig. 4B**). Next, we tested the effect of spiking the sample with 6 mM SDS which resulted in diverse effects on the oligomer sample. The original sample which was prepared in low ionic strength buffer and aged for 24 days post-SEC, displayed mostly connected strings and few single globular particles before SDS addition (**Fig. 4A**). SDS addition resulted in the breakdown of strings and aggregated patches, yielding a more homogeneous sample composed of shorter strings (**Fig. 4C**). Conversely, another sample, prepared in low ionic strength buffer and aged for 36 days post-SEC (**Fig. S2-A**), exhibited non-uniform distribution on the EM grid post SDS addition. Some areas showed a more homogeneous distribution of short strings and globular particles (**Fig. S2-B**), while others contained large patches of aggregated particles (**Fig. S2-C**). Finally, we investigated the effect of adding Aβ42 monomers to 6 days old samples that were primarily composed of strings (**Fig. 4D**). Contrary to previous studies on fibrils, where addition of monomers elongated the fibrils, addition of monomers to strings caused immediate degradation of the strings (**Fig. 4E**)

**Fig. 4.**
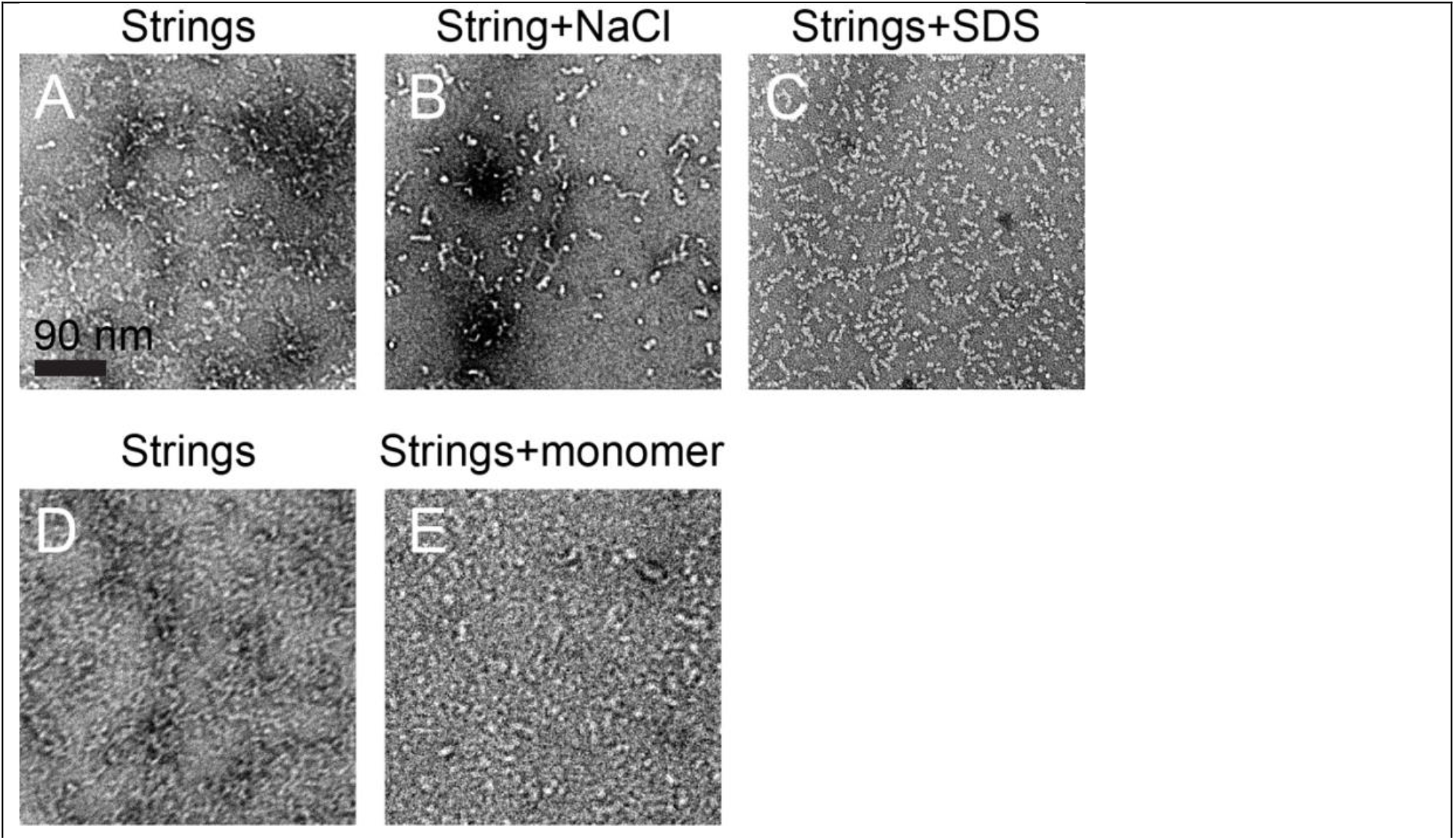
The effects of salt, SDS, and Aβ42 monomer on Aβ42 oligomer sample. A) Aβ42 oligomer sample imaged after 24 days following SEC. Most of the sample consisted of strings. B) An image of the same samples as in Panel A, but after adding NaCl to increase the NaCl concentration to 6 mM. We observed shorter strings and more individual oligomers. C) An image of the same sample as in Panel A, but SDS added to increase the SDS concentration to ∼6 mM. We observed a shorter, more homogeneous string population with some single oligomers. D) Aβ42 oligomer sample imaged 6 days after SEC, showing a high number of strings. E) The same sample as in Panel D, imaged after addition of monomer to it (final Aβ42 monomer concentration ∼89 μM), showing string’s dissociation.

### Cryo-EM on Aβ42 strings suggests that they are associated Aβ42 oligomers with pores perpendicular to the long axis

Here, we investigated the structure of the strings using single particle cryo-EM (**Fig. 5A**). In our previous studies, cryo-EM of 150 kDa Aβ42 oligomers revealed that they resemble globular particles with four-fold symmetry and a central pore (45). Initially, 2,196,587 string particles were picked and were subjected to 2D classification. Several rounds of classification and selection were performed. The resulting 2D class averages (**Fig. 5B**). (obtained from 94,115 string particles) showed that the strings appear to be assemblies of globular 150 kDa Aβ42 oligomers with a ∼16 Å hole in the center that is consistent with the central hole previously observed in the 150 kDa Aβ42 oligomers. Notably, there is not a consistent oligomerization pattern; the strings do not appear to possess helical symmetry. In this way, the strings are different from the classical Aβ42 fibrils, which are helical assemblies of Aβ42 monomers. Most class averages showed clear densities for oligomers of 2-4 globular particles with blurry densities at either end, which is consistent with an incoherent linear assembly. Inspection of the cryo-EM micrographs revealed that the strings appear to be assembled from 3 or more globular particles on average and ranged in size from two globular particles to dozens in length.

**Fig. 5.**
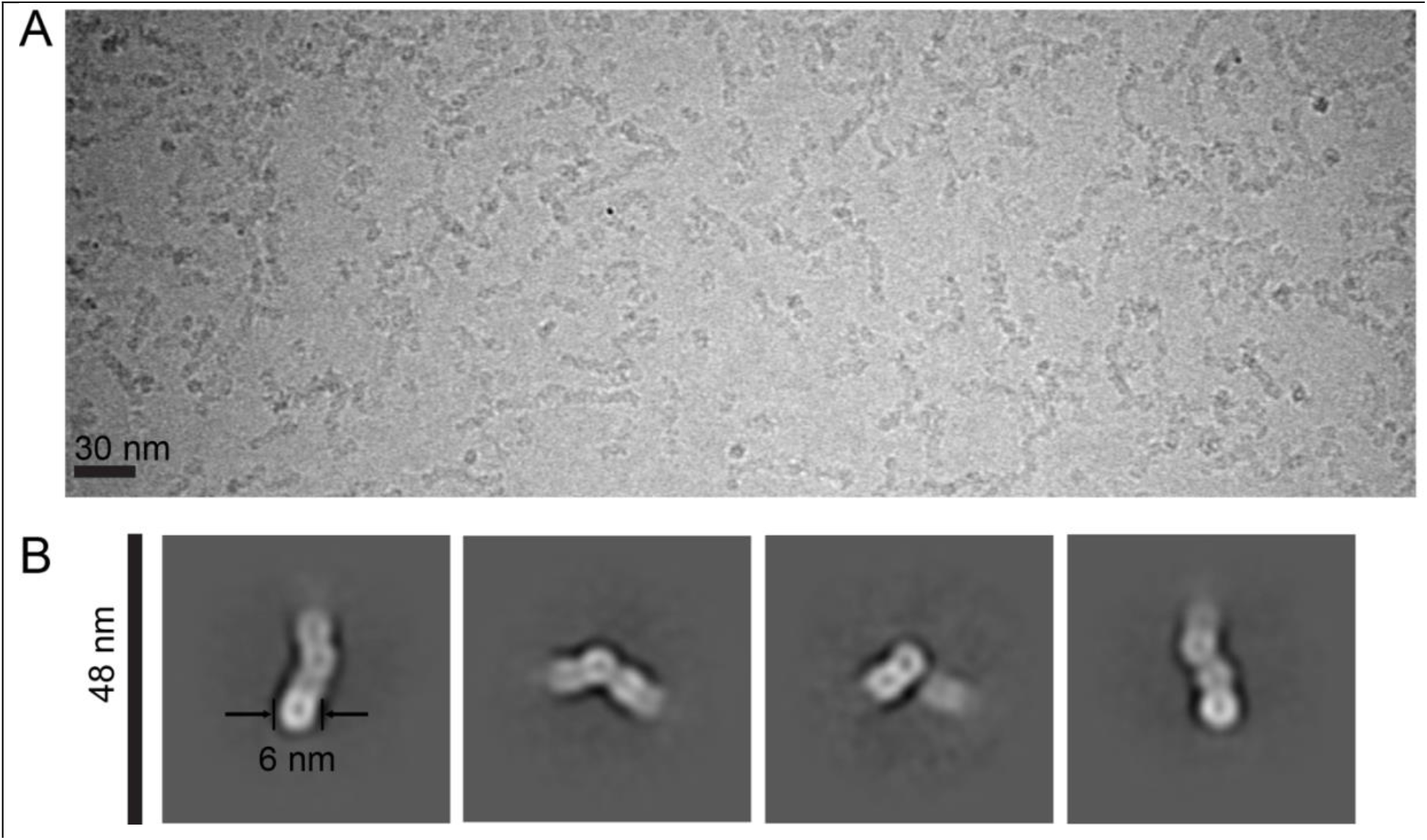
Cryo-EM reveals that Aβ42 strings are laterally associated Aβ42 oligomers. A) Representative micrograph depicting the presence of strings in 150 kDa Aβ42 oligomer sample. B) 2D class averages of strings.

## V. Discussion

We evaluated the temporal dynamics of aggregation of the 150 kDa Aβ42 oligomer, revealing previously uncharacterized aspects of non-fibrillar aggregation. As previously reported, the 150 kDa Aβ42 oligomer is considered “off-pathway” to fibril formation (46,47). In order words, the 150 kDa Aβ42 oligomer does not exhibit a tendency to assemble into thioflavin-positive fibrils or seed fibril growth in monomer solutions (41). Notably, 150 kDa Aβ42 oligomers are not composed of in-register parallel β-sheets, a common structural motif for Aβ amyloid fibrils (15). Nevertheless, our results presented here support the interpretation that the 150 kDa Aβ42 oligomer undergoes a form of self-assembly that begins immediately after oligomer isolation. This self-association represents a non-fibrillar mode of aggregation in the 150 kDa Aβ42 oligomer sample. We use the term “strings” to describe the elongated aspect ratio of this self-assembly while differentiating these structures from fibrils. Although this self-association is progressive over time, the length of the strings does not appear to exceed a few hundred nanometers. To summarize our experimental observations, TEM revealed the elongated nature of strings and the appearance of subunits similar in size to 150 kDa Aβ42 oligomers, while SEC and DLS confirmed the increase in particle size distributions in bulk solution with time. Analysis of the string’s length showed significant variability, with sizes ranging from tens of nanometers to a few hundred nanometers. However, we did not observe strings extending as long as a micrometer, highlighting the difference between strings and amyloid fibrils.

We also found string formation to be sensitive to ionic strength, SDS addition, and Aβ42 monomer addition, revealing insights into oligomer-oligomer interactions. We prepared oligomer samples in two different ionic strength conditions. High ionic strength corresponded to slower string elongation accompanied by the appearance of doughnut-shaped structures. We suggest that doughnuts result from short strings doubling back on themselves to form rings. In other words, we hypothesize that doughnuts and strings have the same underlying molecular structure, as doughnut wall thickness (∼6 nm) appears to be the same as string widths and 150 kDa Aβ42 oligomer diameters. We also demonstrated that string formation is reversible; string dissociation can be induced by adding NaCl, SDS, or Aβ42 monomers. These results suggest that electrostatic interactions, SDS detergent, and interactions with Aβ42 monomers readily affect the equilibrium required for string formation. We also observed that SDS affects the morphology of the aggregated string assemblies. The results of Aβ42 monomer addition to strings is in contrast to the behavior of amyloid fibrils: while fibrils can have string-like lengths (e.g., when fragmented via sonication), they would grow rather than dissociate upon Aβ monomer addition. It is presently unclear why strings and fibrils behave differently with Aβ monomer addition, but we suggest that the difference may be related to the string formation via the assembly of 150 kDa Aβ42 oligomers compared to fibril elongation by monomer addition. If the strings were to grow by monomer addition, the addition of more monomer would not have impacted them or may have even caused further growth. Therefore, 150 kDa Aβ42 oligomers exhibit a different type of assembly mediated by the self-association of larger-than-monomeric particles (a schematic model of our observations is presented in **Fig. S3**). Oligomer self-association does not appear to be mediated by highly ordered molecular interactions as with fibrils, which may explain why the size of strings is limited to much less than a micrometer. It is notable that while we consider 150 kDa Aβ42 oligomer and string formation to be products of a distinct aggregation pathway from fibril formation, 150 kDa Aβ42 oligomers and strings might eventually convert into fibrils if they first dissociate in dynamic equilibrium with monomers.

Cryo-EM analysis confirms that strings exhibit a configuration resembling associated 150 kDa Aβ42 oligomeric particles. Specifically, our 2D classification results on strings exhibited subunits with the same size as 150 kDa Aβ42 oligomers and central pores resembling the pores we previously observed in 150 kDa Aβ42 oligomers. The oligomers are arranged such that the pores are oriented perpendicular to the long axis of each string. In our effort to model the molecular structure of the 150 kDa Aβ42 oligomers, we proposed that the pore is oriented parallel to the β-sheet axis, in contrast to the structures of Aβ fibrils. To be clear, while fibril β-sheets extend in the direction of the long axis of each fibril, our present results suggest that β-sheets within strings are perpendicular to the long axis of each string. This distinction is consistent with the differences we observed between strings and fibrils, potentially explaining why strings fail to elongate to typical fibrillar dimensions (fibrils lengths are microns and above).

Finally, we suggest that our presented data highlights the important challenges to the field of Aβ pathological aggregation. While research has underscored the potentially pivotal role of Aβ oligomers in the pathogenesis of AD, we do not possess detailed structural knowledge of oligomer aggregation and how oligomer formation differs from fibril formation. Small size implies higher diffusivity and more interactions with neuronal membranes. It has been suggested that oligomers can create pores in neuronal cell membranes. While these characteristics have been associated with disease, our current understanding remains limited to their small size and predominant β-sheet structure. We desire molecular-level knowledge of pathological mechanisms, such as proposed neuronal membrane disruption, which would benefit from the knowledge of oligomer and string assembly pathways. A detailed high-resolution three-dimensional structure of the 150 kDa Aβ42 oligomer would be of great interest. Moreover, this structural knowledge could provide valuable cues for the development of targeted therapeutic interventions aimed at mitigating the aggregation and deleterious effects of Aβ, potentially fostering the creation of more efficacious treatments for AD and other neurodegenerative ailments.

## ACKNOWLEDGEMENTS

This work was supported by the National Institute of Health (RF1AG073434). Cryo-EM data collection was performed at New York Structural Biology Center (NCCAT). We gratefully acknowledge Hui Wei for conducting the data collection and assistance with sample preparation for cryo-EM. We thank Dr. Peter Randolph at the Institute of Molecular Biophysics at FSU for his assistance with DLS experiments.

## Author Contributions

J.W prepared the monomer samples. S.K prepared the oligomer samples. S.K and S.S designed the experiments. S.K performed negative stain TEM visualizations, SEC and DLS experiments. S.K performed cryo-EM analysis with S.S supervision. S.K prepared the figures. S.K, A.P and S.S wrote the manuscript. T.R, A.P and S.S conceptualized, administrated and supervised the project. All authors read and edited the manuscript.

