## Supplementary Material for "Aggregation Dynamics of a 150 kDa Aβ42 Oligomer: Insights from Cryo Electron Microscopy and Multimodal Analysis"

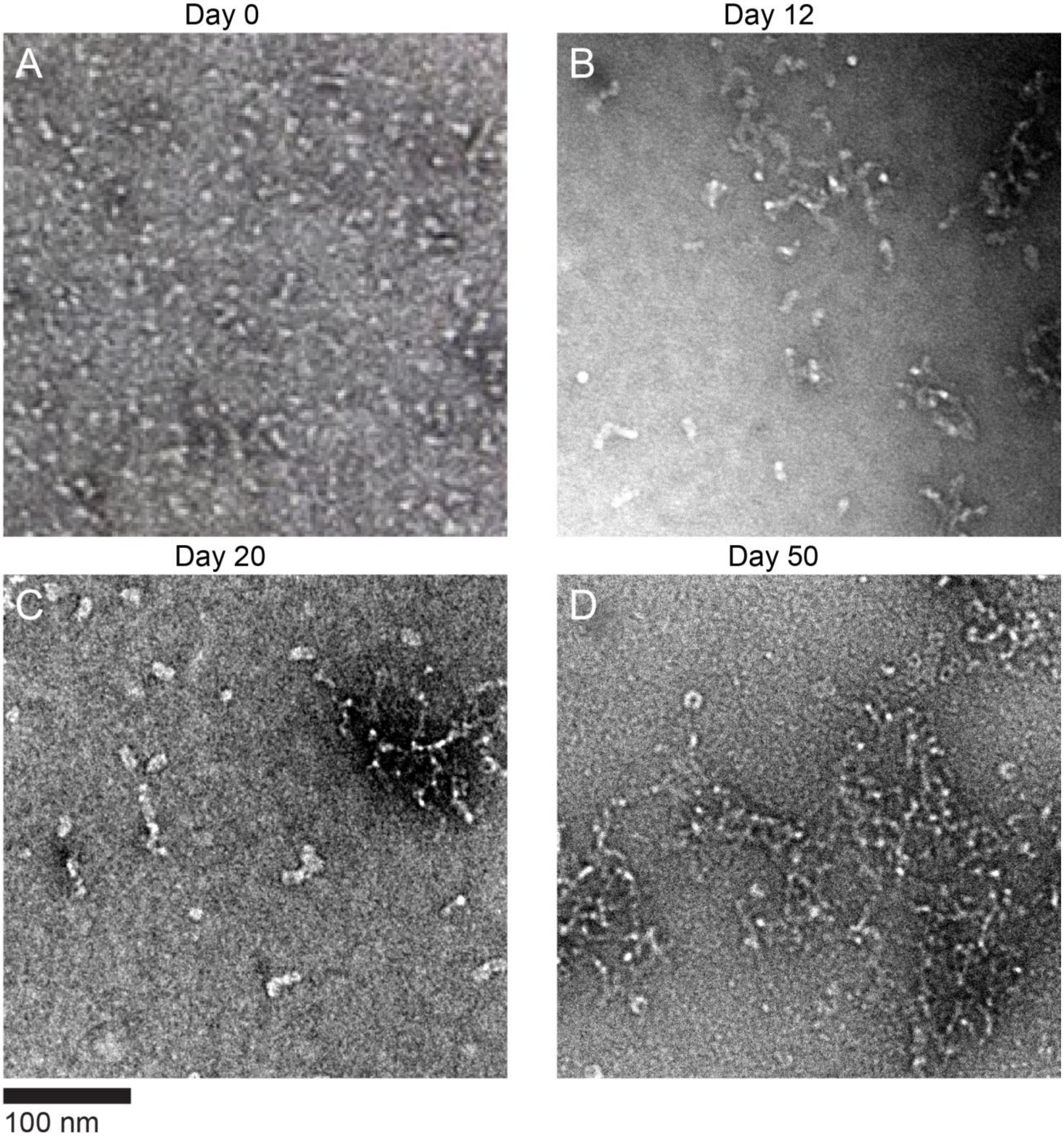


**Fig. S1.** Time course of negative stain TEM images of the same Aβ42 oligomer sample prepared in high ionic strength buffer (20 mM sodium phosphate pH 7.5 plus 50 mM NaCl). String population/length appears to increase over time following SEC isolation of 150 kDa Aβ42 oligomers. A) Sample imaged immediately after SEC. B) Sample imaged 12 after SEC. C) Sample imaged 20 days after SEC. D) Sample imaged 50 days after SEC.

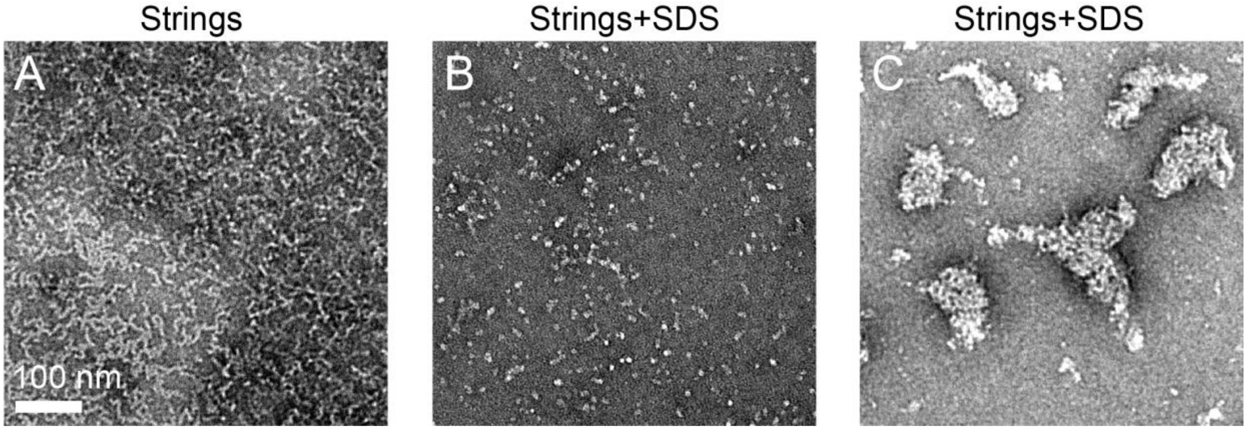


**Fig. S2.** The effects of SDS on Aβ42 oligomer sample prepared in high ionic strength buffer containing 20 mM sodium phosphate pH 7.5 plus 50 mM NaCl. A) Aβ42 oligomer sample imaged after 36 days following SEC. Most of the sample consisted of strings. B) An image of the same samples as in Panel A, but after adding SDS to increase the SDS concentration to ~6 mM. We observed shorter strings and more individual oligomers. C) An image of the same sample as in Panel B, but in another region of the grid. We observed cloud shape patches of strings.

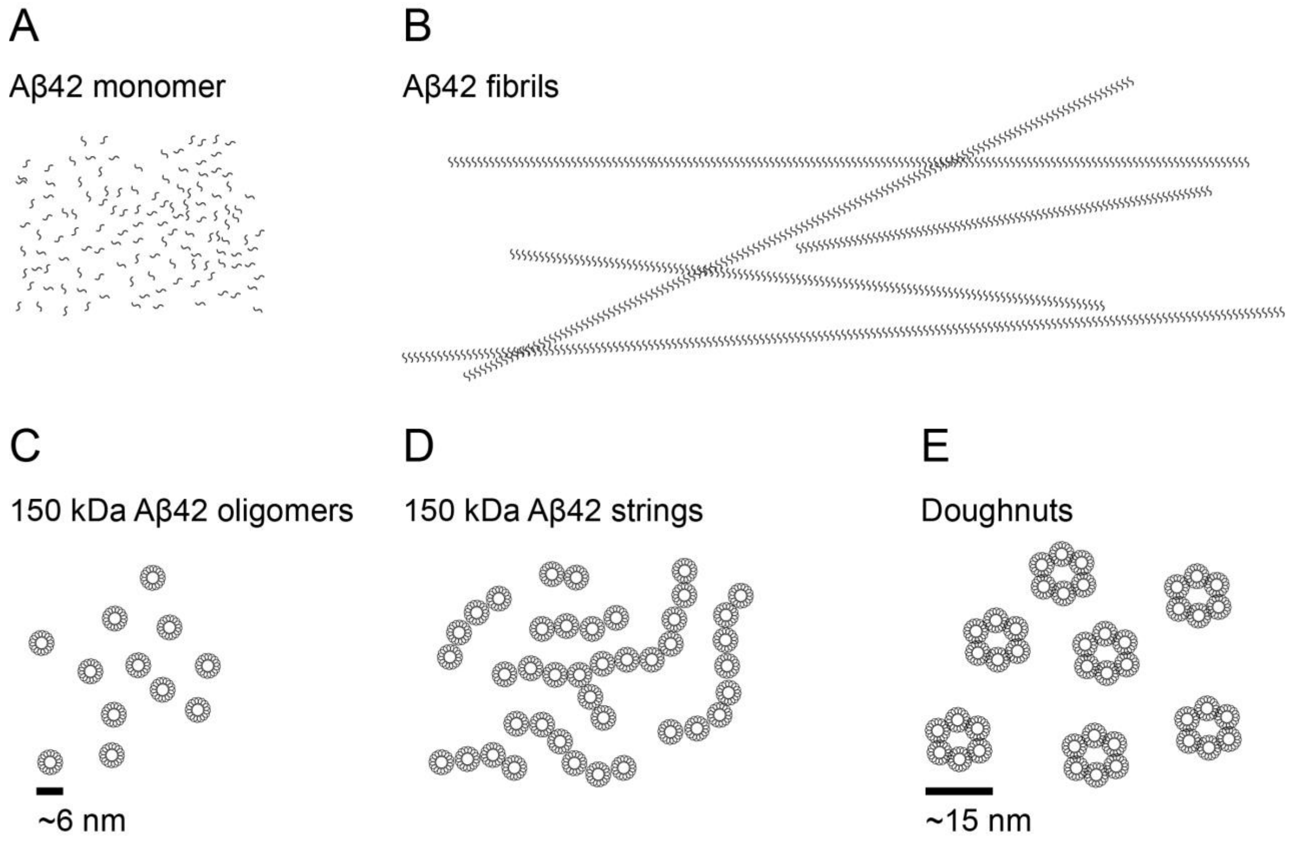


**Fig. S3.** Schematic model representation of Aβ42 monomer, classic fibrils, 150 kDa oligomers, strings and doughnuts. A) Schematic representation of Aβ42 monomers. B) Schematic representation of classic Aβ42 fibrils, fibrils can grow in length to several microns. C) Schematic representation of classic 150 kDa Aβ42 oligomers D) Schematic representation of Aβ42 strings. Strings vary in length, but they are limited in size to few hundred nano meters. E) Doughnut shape entities occasionally observed in Aβ42 oligomer sample. They are more prevalent in samples prepared in higher ionic strength condition compared to the samples prepared in lower ionic strength.
